# Single-dose DMT reverses anhedonia and cognitive deficits via restoration of neurogenesis in a stress-induced depression model

**DOI:** 10.1101/2025.04.26.650765

**Authors:** Rafael V. Lima da Cruz, Rêmullo B. G. de Miranda Costa, Gabriel M. de Queiroz, Tijana Stojanovic, Thiago C. Moulin, Richardson N. Leão

## Abstract

Major depressive disorder (MDD) remains a leading cause of disability worldwide, with current treatments limited by delayed onset and low efficacy. The serotonergic psychedelic N,N-dimethyltryptamine (DMT) has shown rapid antidepressant effects in early clinical studies, yet its mechanisms and efficacy remain poorly characterized in established models of depression. Here, we evaluated the effects of a single dose of DMT in the Chronic Unpredictable Mild Stress (UCMS) paradigm, a robust mouse model recapitulating key features of MDD, including anhedonia and cognitive impairment. DMT administered after UCMS reversed depressive-like behavior and restored cognitive performance, outperforming chronic fluoxetine across most domains. When administered during the stress period, DMT prevented the development of anhedonia but did not rescue cognitive deficits, suggesting partial protection. Notably, DMT remained effective under isoflurane anesthesia, indicating that its therapeutic action can occur independently of the psychedelic experience. Histological analyses revealed that all DMT regimes significantly increased adult-born granule cell (abGC) integration and reduced the number of ectopically abnormally integrated abGCs,. Together, our findings highlight the robust and multifaceted effects of DMT on behavior and neurogenesis, positioning it as a promising candidate for rapid-acting antidepressant strategies that target structural circuit repair.

## Introduction

Major depressive disorder (MDD) is among the most prevalent and debilitating mental illnesses worldwide. According to recent estimates, up to 20% of individuals in developed countries may experience at least one major depressive episode in their lifetime [WHO; 2020]. This disorder is strongly associated with reduced quality of life, cognitive abilities, and work productivity [1]. While conventional antidepressants such as selective serotonin reuptake inhibitors (SSRIs) have demonstrated therapeutic efficacy, their clinical limitations are substantial. These include a delayed onset of therapeutic action, often taking several weeks, and a high rate of treatment resistance, with approximately 30% of patients showing inadequate responses despite optimized interventions [2]. This has created a critical need for the development of more effective and fast-acting treatment strategies.

In recent years, accumulating evidence has highlighted the role of adult hippocampal neurogenesis in the pathophysiology and treatment of depression. The dentate gyrus (DG), a subregion of the hippocampus, continually integrates adult-born granule cells (abGCs) into its circuitry, a process essential for cognitive flexibility and mood regulation [3, 4]. Activation of abGCs has been shown to be sufficient to alleviate depression-like phenotypes, reversing the effects of chronic stress [5]. Additionally, most effective antidepressant treatments associated with improved behavioral outcomes enhance neurogenesis, including SSRIs [6, 7], lithium [8, 9] ketamine [10, 11] and even non-pharmacological interventions such as electroconvulsive shock [12, 13]. These findings support the neurogenic hypothesis of depression, which posits that disrupted neurogenesis contributes to depressive symptomatology and, thus, targeting of neurogenic factors may yield therapeutic benefits [3, 14].

Among emerging therapeutic options, serotonergic psychedelics have garnered significant attention for their rapid and sustained antidepressant effects. Compounds such as psilocybin, LSD, and mescaline, classified as classical psychedelics due to their potent 5-HT2A receptor agonism, have demonstrated the ability to rapidly alleviate depressive symptoms, often within hours after a single administration [15–18]. Psilocybin, for example, has shown robust and prolonged antidepressant efficacy in clinical trials, with symptom improvements persisting for up to 12 months in some patients, and a favourable safety profile, lacking addictive potential and eliciting no serious adverse effects [19–23]. Notably, these compounds have been shown to enhance neuroplasticity and promote various aspects of neurogenesis, including the proliferation, migration, and differentiation of neurons [24]. ‘N,N-dimethyltryptamine (DMT) is another promising antidepressant compound. It is a key psychoactive ingredient in ayahuasca, a traditional Amazonian brew used ceremonially for centuries25. The long-standing, widespread use of DMT across various populations for centuries bolsters the evidence for its potential safety profile in the context of antidepressant applications with remarkable tolerability [25]. Like psilocybin, DMT is a serotonergic hallucinogen with slightly higher affinity for 5-HT2A receptors. Furthermore, the DMT molecule naturally occurs in various organisms, including humans [26]. When administered intravenously or inhaled, DMT bypasses rapid first-pass metabolism and induces intense but short-lived psychedelic experiences, characterized by profound alterations in perception, affect, and cognition [27, 28]. Despite decades of observational and pharmacological research, the clinical potential of DMT has only recently begun to be systematically evaluated [29]. For example, Osorio et al. [30] reported rapid reductions in depression scores following a single dose of ayahuasca in treatment-resistant patients, with effects sustained over several weeks. Palhano-Fontes et al. [31] conducted a randomized placebo-controlled trial confirming that ayahuasca produced a significant and rapid antidepressant effect compared to placebo. In a longitudinal study, Jiménez-Garrido et al. [32] reported that over 80% of participants who met the diagnostic criteria for a psychiatric disorder exhibited sustained clinical improvements six months after ayahuasca use. More recently, studies administering pure DMT have begun to emerge. For instance, A recent open-label trial showed that inhaled DMT produced rapid and sustained antidepressant effects in patients with treatment-resistant depression [33]. The treatment was safe, well-tolerated, and led to a significant reduction in depressive symptoms within 7 days, with response and remission rates of 85.7% and 57.1%, respectively. Effects lasted up to three months, and suicidal ideation dropped significantly, highlighting DMT’s potential as a fast-acting and scalable therapeutic option.,. Moreover, in an exploratory open-label trial for dose-related effects, intravenous DMT administration to MDD treatment-resistant individuals produced a rapid and significant decrease in depression scores, with mostly mild adverse events reported [34].

Beyond psychiatric outcomes, recent preclinical studies have elucidated DMT’s ability to modulate neural plasticity. Rodent models have shown that both high-dose and microdosed DMT facilitate fear-extinction learning and produce antidepressant-like effects [35, 36]. Likewise, 5-MeO-DMT, a structural analog of DMT with similar pharmacodynamic properties, has shown robust anxiolytic effects in rodents subjected to stress [37, 38]. At the cellular level, DMT enhances neurite outgrowth, synaptic density, and dendritic spine formation in cortical neurons, comparable to the effects of ketamine [39]. Moreover, our group has previously shown that a single 5-MeO-DMT administration increases adult hippocampal neurogenesis and alters the electrophysiological properties of newborn neurons in the DG, conferring heightened excitability and synaptic plasticity to abGCs [40]. These neuroplastic changes are further supported by molecular findings showing upregulation of brain-derived neurotrophic factor (BDNF) and activation of sigma-1 receptors, both implicated in mood regulation and neurogenesis [24, 41, 42].

Despite promising clinical and preclinical findings, key questions remain regarding the therapeutic potential of DMT in depression. Notably, no study to date has systematically examined the effects of pure DMT in a validated preclinical model of depression-like behavior. Here, we address this critical gap by using the Unpredictable Chronic Mild Stress (UCMS) paradigm in mice, a well-established model that captures core features of MDD, particularly anhedonia [43]. Furthermore, while previous studies have shown that DMT can reverse depressive symptoms, it remains unknown whether it can prevent their onset when administered during periods of chronic stress. To investigate this, we administered a single dose of DMT not only after, but also during the UCMS protocol. A final unresolved issue concerns the role of subjective psychedelic experiences in mediating therapeutic effects. While some authors argue that the hallucinogenic conscious experience is necessary for enduring clinical benefits [44], others suggest that psychedelics act through downstream molecular and synaptic mechanisms that might be separable from conscious perception43. We tested this by administering DMT under isoflurane anesthesia, thereby suppressing the acute wake experience. We compared the effects of these different DMT treatment regimens to saline and fluoxetine controls, assessing behavioral outcomes across depressive, anxiogenic, and cognitive domains post-UCMS. We also performed histological analysis of abGCs, which were temporally labeled using the DCXCreERT2::TdTOMlox/lox system. Together, our findings offer novel phenotypic and mechanistic insights into how altered neurogenesis and abGC circuit integration contribute to the pathophysiology of depression and its reversal by psychedelics such as DMT.

## Results

### Depression-like phenotypes

Mice were exposed to 56 days of UCMS before receiving a single injection of either DMT, to test if acute psychedelic treatment is able to reverse UCMS-induced phenotypes; DMT under isoflurane anesthesia (DMT+Iso), to examine the effects of conscious psychedelic experience; or vehicle control (saline). A parallel cohort DMT during the stress period (DMT day 28) to assess whether early intervention can prevent the development of stress-induced phenotypes. Finally, a positive control group was treated with daily fluoxetine, a conventional antidepressant, for 30 days partially overlapping with the UCMS protocol (Figure 1A).

**Figure 1.**
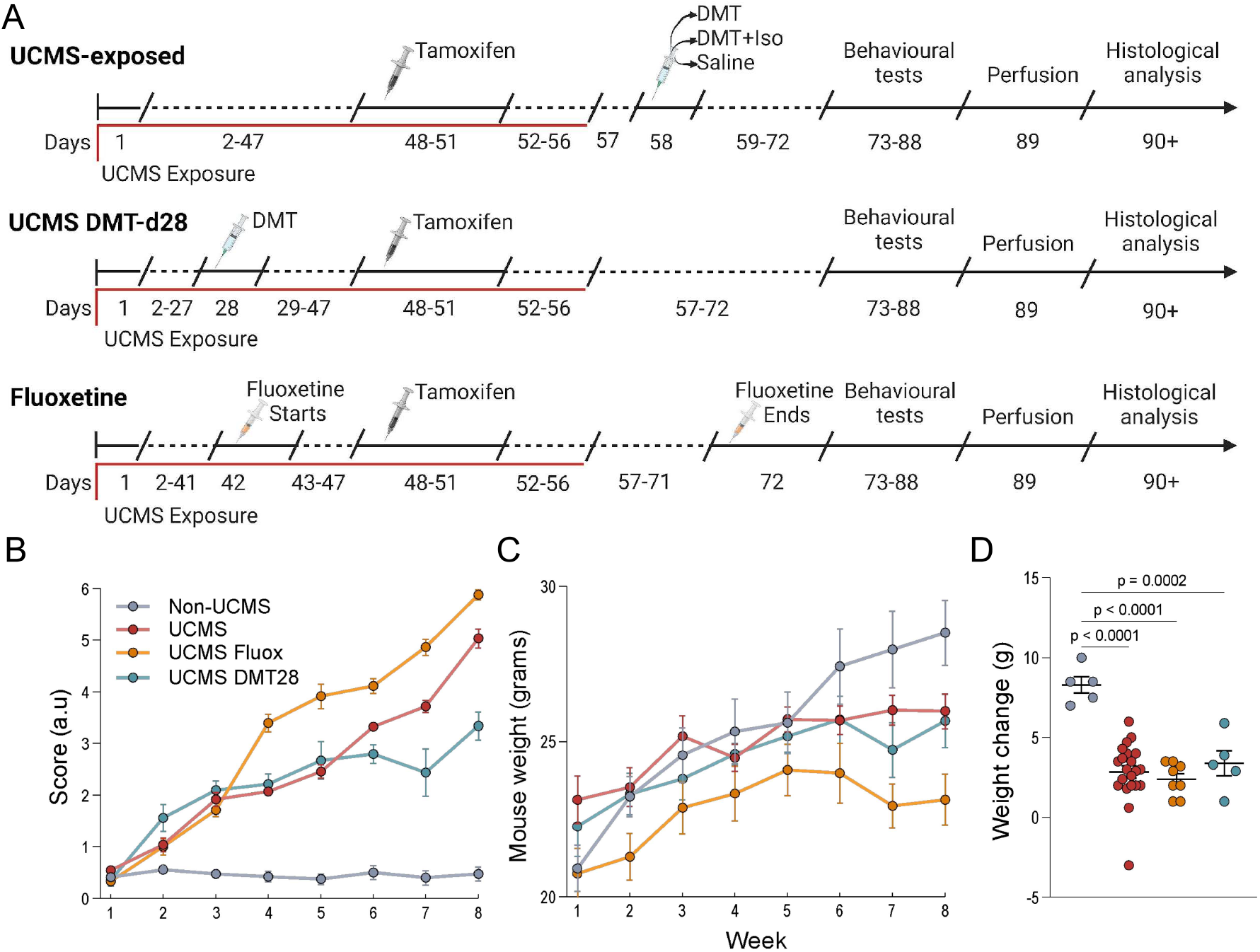
Experimental Design and Outcome Measures. (A) Timeline of experimental groups. The upper row represents UCMS-exposed animals, which underwent the UCMS paradigm from days 1–47, followed by a single injection of either Saline, DMT, or DMT+Iso on day 58. The middle row represents the UCMS DMT-d28 group, which followed the same UCMS exposure timeline but received a single DMT treatment on day 28. The bottom row represents the Fluoxetine group, in which animals started chronic fluoxetine treatment on day 42, continuing through day 72, overlapping with UCMS exposure. All groups received tamoxifen (days 48–51) and underwent behavioral testing (days 73–88), perfusion (day 89), and histological analysis (days 90+). (B) Longitudinal coat state scoring, where higher scores indicate worsening fur condition. (C) Longitudinal body weight assessment. (D) Comparison of final weight gain (week 8 – week 1) across experimental groups. Analyses in panels were conducted using either two-way ANOVA (in B and C) or one-way ANOVA followed by Tukey’s post-hoc test (in D); adjusted p-values are shown in the figure. Sample sizes: Non-UCMS (n = 5), UCMS (n = 21), UCMS Fluoxetine (n = 8), and UCMS DMT-d28 (n = 5).

Consistent with previous findings, weekly monitoring revealed that UCMS-exposed mice had significantly deteriorated coat state and reduced weight gain compared to Non-UCMS controls (Figure 1B-D), both established indicators of stress-induced behavioral alterations 44. Specifically, two-way ANOVA of coat scores indicated significant main effects of time, treatment, and their interaction (all p < 0.0001). Post-hoc analysis (Holm-Sidak multiple comparison test) at the final measurement confirmed that all UCMS groups showed significantly worse coat conditions than Non-UCMS controls (p < 0.0001 for UCMS and UCMS Fluoxetine; p = 0.0004 for UCMS DMT-d28). Interestingly, mice receiving DMT midway through the stress protocol (UCMS DMT-d28) showed a modest yet significant improvement in coat state compared to both UCMS alone (p = 0.0042) and UCMS Fluoxetine groups (p = 0.0012). Notably, fluoxetine-treated UCMS mice displayed the worst coat condition among stressed groups, significantly worse even compared to UCMS alone (p = 0.0020). Moreover, two-way ANOVA analysis of weight gain showed significant effects of time and the interaction between time and treatment (both p < 0.0001), but no main effect of treatment alone. To clarify these differences, a subsequent one-way ANOVA was conducted on the total weight change (Δ weight) during the UCMS protocol (Figure 1F), revealing a significant difference among groups (p < 0.0001). Post-hoc analysis (Tukey’s multiple comparisons test) demonstrated that all UCMS groups experienced significantly reduced weight gain compared to Non-UCMS controls (Non-UCMS vs. UCMS, p < 0.0001; Non-UCMS vs. UCMS Fluoxetine, p < 0.0001; Non-UCMS vs. UCMS DMT-d28, p = 0.0002). However, no significant differences were found among the UCMS groups themselves (UCMS vs. UCMS Fluoxetine, p = 0.8992; UCMS vs. UCMS DMT-d28, p = 0.9083; UCMS Fluoxetine vs. UCMS DMT-d28, p = 0.6990), indicating that this protocol had consistent effects across groups.

Then, to evaluate reward-seeking behavior, sucrose preference tests (SPTs) were conducted before and after UCMS exposure, as well as after treatments (Figure 2A). To first assess the impact of UCMS itself, STPs were compared across the stress-exposed cohorts prior to any treatment (Figure 2B). Then, one-way ANOVA analyses followed by Tukey’s multiple comparisons test of sucrose preference change between the two assessments (Figure 2C). As expected, untreated UCMS-exposed mice exhibited a marked reduction in sucrose preference when compared to Non-UCMS controls (p < 0.0001), indicating a robust anhedonic response. Notably, mice in the UCMS DMT-d28 group, treated during the stress period, showed significantly less anhedonia in relation to the untreated UCMS group (p < 0.0001), and were comparable to Non-UCMS controls (p = 0.3723). In contrast, initiating fluoxetine treatment during the later stages of UCMS did not prevent the development of anhedonia, as sucrose preference remained significantly lower than Non-UCMS levels (p = 0.0044) and similar to that of the untreated UCMS group (p = 0.4933).

**Figure 2.**
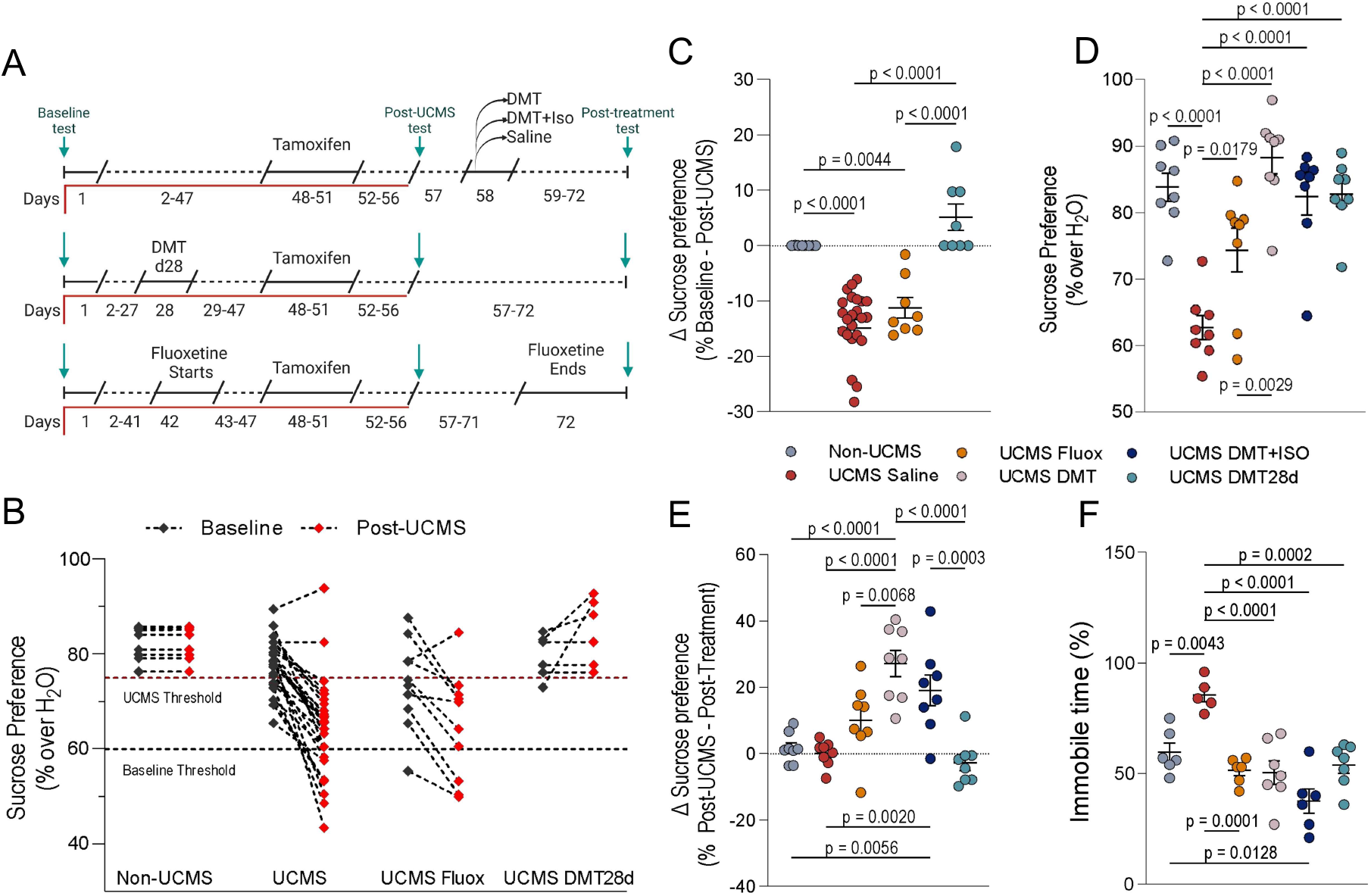
DMT and fluoxetine treatments differentially reverse UCMS-induced anhedonia and behavioral despair. (A) Experimental timeline highlighting behavioral testing and treatment periods. Green arrows indicate behavioral test timepoints. Sucrose preference tests were conducted at baseline, after UCMS exposure (pre-treatment), and after treatment across all groups. Tail suspension test was performed only at the post-treatment timepoint. Treatment regimens are the same as shown in Figure 1A: acute DMT, DMT+Iso, or Saline on day 58; DMT-d28 during the UCMS protocol; and daily fluoxetine from days 42–71. (B) Sucrose preference before and after UCMS exposure, prior to treatment, illustrating stress-induced anhedonia across groups. A black dashed line marks the 60% baseline exclusion threshold; mice showing unusually low sucrose preference before UCMS (n = 1) were considered sucrose non-responders and excluded. A red dashed line at 75% indicates the post-UCMS exclusion threshold; mice that maintained high sucrose preference after UCMS without treatment (n = 2) were excluded from further intervention due to stress insensitivity. (C) Change in sucrose preference (Δ sucrose preference) from baseline to post-UCMS for each group shown in panel B. (D) Post-treatment sucrose preference across all groups. (E) Change in sucrose preference (Δ sucrose preference) from pre-to post-treatment. (F) Tail suspension test immobility time measured after treatment to assess behavioral despair. All data are presented as individual values with mean ± SEM (panels C–F); panel B shows individual values only. Analyses in panels C–F were conducted using one-way ANOVA followed by Tukey’s post-hoc test; adjusted p-values are shown in the figure. Sample sizes: (B, after exclusions): Non-UCMS (n = 8), UCMS (n = 21), UCMS Fluoxetine (n = 8), UCMS DMT-d28 (n = 5). (C): same as (B). (D–E): all groups n = 8. (F): Non-UCMS (n = 6), UCMS Saline (n = 5), UCMS Fluoxetine (n = 6), UCMS DMT (n = 7), UCMS DMT+Iso (n = 6), UCMS DMT-d28 (n = 7).

We then assessed changes in sucrose preference between the end of the Since this test requires frame-by-frame pose estimation, we used DeepLabCut for tracking. A custom Python script was developed to analyze the DLC output controls (p = 0.0043), reflecting the development of a depressive-like state. Remarkably, all tested treatments, i.e., DMT (p < 0.0001), DMT+Iso (p < 0.0001), fluoxetine (p = 0.0001), and DMT-d28 (p = 0.0002), effectively reduced immobility compared to UCMS saline controls. Despite targeting different timepoints and mechanisms, these interventions produced similarly robust effects in reversing helplessness behavior. No significant differences were detected among the treatment groups themselves (all p > 0.09), suggesting convergent efficacy in this assay. Taken together with the sucrose preference data, these results support a broad antidepressant-like action of DMT and highlight its potential to counteract distinct dimensions of depression-related behavior.

### Anxiety-like phenotypes

In the OFT (Supplementary Figure 1A–C), we did not observe any statistically significant differences between UCMS saline-treated mice, Non-UCMS controls, or any of the treatment groups following one-way ANOVA analysis of the distance travelled. Similarly, no significant differences were found among groups when measuring time spent in the center, expressed as a percentage of total test duration. Representative tracking paths and occupancy heatmaps from a sample animal are shown in Supplementary Figure 1A and illustrate the spatial distribution of exploration across the arena.

Likewise, in the EPM (Supplementary Figure 1D–F), ANOVA analysis revealed no significant differences across treatment groups, UCMS saline, or Non-UCMS controls in terms of the percentage of time spent in the open arms. We also found no significant differences in the number of head dips, an index of exploratory behavior. A representative example of the movement trajectory in the EPM is shown in Supplementary Figure 1D, highlighting typical arm entries and tracking across arms. Taken together, these results suggest that neither UCMS nor any of the pharmacological treatments tested induced measurable changes in anxiety-like behavior under the current experimental conditions.

### Cognitive Phenotypes

To assess cognitive function and specifically examine pattern separation, we employed the DNMP-RAM task, illustrated in Figure 3A. Performance analysis for lower difficulty, longer separation conditions (S3 and S4) revealed no significant differences across groups (Figure 3B, right panel). However, significant group differences emerged at the higher difficulty, lower separation level (S2) of the task (Figure 3B, left panel). Specifically, ANOVA followed by Tukey’s multiple comparisons test showed that UCMS-saline mice performed significantly worse than naïve controls (Non-UCMS, p = 0.0165). Moreover, animals treated with fluoxetine (p = 0.0007), DMT (p < 0.0001), and DMT+Iso (p = 0.0108) displayed significantly enhanced performance compared to the UCMS-saline group, indicating a reversal of UCMS-induced cognitive deficits. Conversely, the DMTd28 group did not significantly differ from saline-treated animals (p = 0.7273), indicating no cognitive improvement. We further analysed the learning trajectory across the 15-day period to determine if groups had distinct learning curves during the higher difficulty separation trials (S2). As illustrated in Figure 3C for the higher difficulty task, animals from the UCMS-saline and DMTd28 groups showed minimal to no learning throughout the assessment period, plateauing around the chance-level correct-response rate. Repeated-measures two-way ANOVA analysis showed significant differences between treatments (p<0.0001) and significant effects of training (p<0.0001). Mice treated with fluoxetine, DMT and DMT combined with isoflurane post-UCMS, as well as the Non-UCMS group, exhibited significant higher learning than UCMS-Saline at the 15-day mark (Holm-Sidak multiple-comparisons test; p=0.0001, p=0.0016, p=0.0169, and p=0.0049, respectively). Similar analysis of lower difficulty separation trials (S3 and S4, Figure 3D), revealed a significant effect of training (p<0.0001), but not of treatment (p=0.7677).

**Figure 3.**
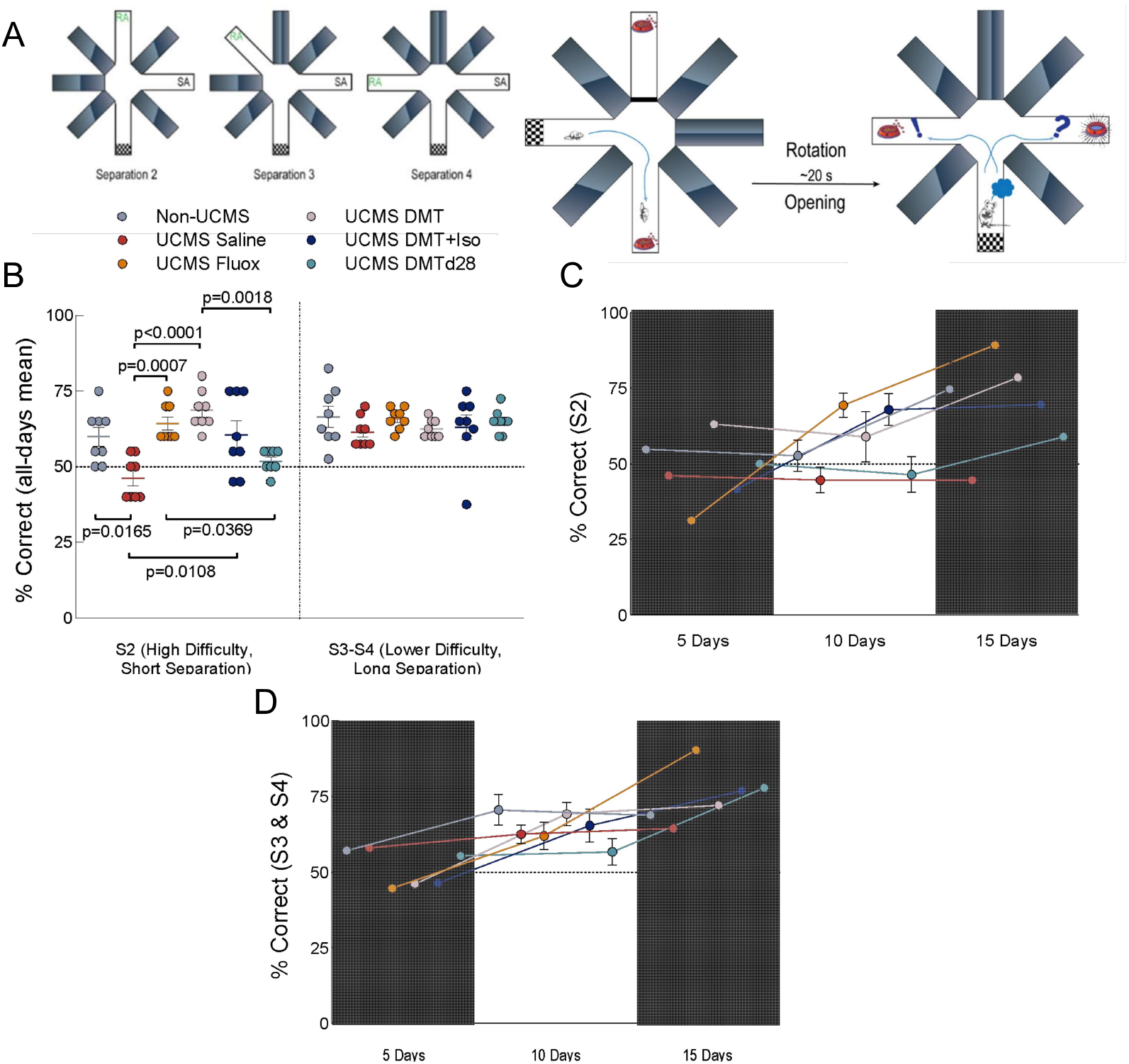
DMT reverses UCMS-induced cognitive impairments. (A) Schematic of the DNMP-RAM task. Left: Task difficulty is manipulated by varying the spatial separation between the sample arm (SA) and the correct choice arm, denoted as S2 (adjacent arm), S3, or S4 (maximally separated). Right: Each trial consists of a sample phase, where the animal is allowed to explore one open arm to retrieve a reward. After a brief delay (∼20s), the maze is rotated, and the animal is returned to the center for the choice phase, where both arms are opened and a reward is delivered only if the animal enters the non-matching arm. (B) Task performance across groups. Left panel: Significant group differences were found for the most difficult condition (S2). UCMS-saline mice showed impaired performance relative to Non-UCMS controls, while animals treated post-UCMS with fluoxetine, DMT, or DMT+Iso performed significantly better than UCMS-saline. Right panel: No significant group differences were detected under lower difficulty (S3 and S4) conditions. (C) Learning trajectories for high-difficulty (S2) trials across the 15-day testing period. Analysis showed significant effects of treatment and training. Non-UCMS, Fluoxetine-, DMT-, and DMT+Iso-treated animals exhibited significantly better performance at the final timepoint compared to UCMS-saline (D) Learning trajectories for lower difficulty (S3/S4) trials. A significant effect of training was observed (p < 0.0001), but no group differences were detected. Analyses in panels were conducted using either one-way ANOVA followed by Tukey’s post-hoc test (in B), or repeated-measures two-way ANOVA with Holm-Sidak post-hoc test (in C and D). All data are presented as individual values with mean ± SEM. Sample size: all groups, n = 8.

### Adult Neurogenesis and Aberrant GC Integration

To evaluate the impact of treatments on GCs neurogenesis and their integration within the DG, we performed confocal microscopy of newborn GCs by the tamoxifen-induced tagging within the genetically modified DCXCreERT2::TdTOMlox/lox mice (Figure 4A, B). Briefly, following behavioral assessments, animals were euthanized, and their brains were collected for detailed histological analysis. DCX-TOM+ cells were systematically quantified along the dorsoventral hippocampal axis, with additional consideration given to ectopically located neurons (i.e., cells integrating outside the granule cell layer, GCL). Histological assessment with one-way ANOVA analyses revealed significant treatment-related differences in the number and distribution of DCX-TOM+ neurons. In the ventral hippocampus (Figure 4C), both the DMT and DMTd28 groups exhibited roughly a two-fold increase in DCX-TOM+ cell counts compared to non-stressed control animals (Non-UCMS vs. DMT p < 0.0001; vs DMTd28 p < 0.0001). These two DMT-treated groups did not differ from one another (p > 0.9999), but each showed significantly greater neurogenesis than fluoxetine-treated mice (p = 0.0076 and p = 0.0066 for DMT and DMT-d28 vs. Fluoxetine, respectively), as well as those receiving DMT under isoflurane (p = 0.0483 and p = 0.0431). DMT+Iso was still significantly more effective than UCMS-saline (p = 0.0245), while fluoxetine treatment did not have a significant effect (p = 0.1288).

**Figure 4.**
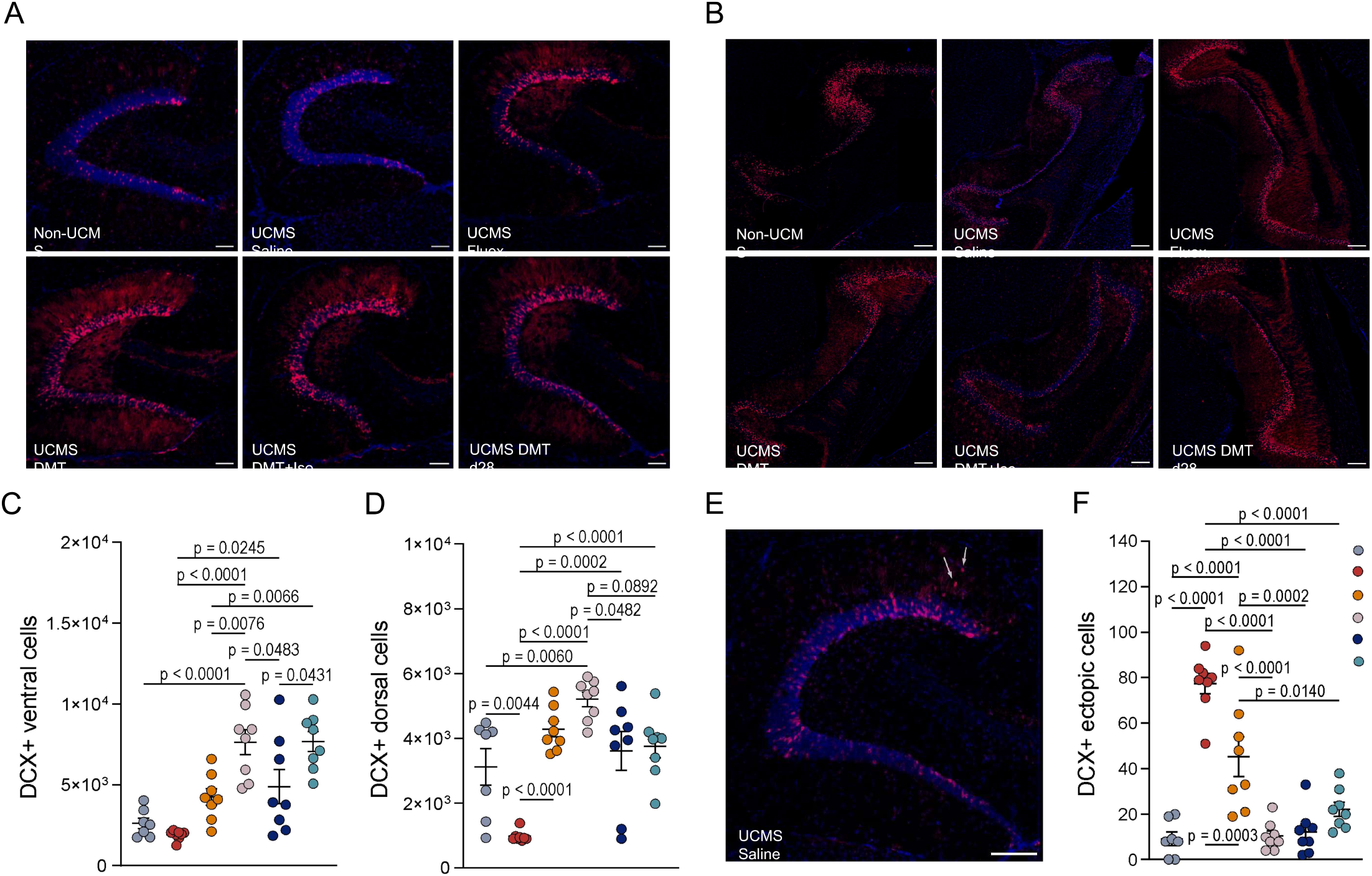
DMT promotes adult hippocampal neurogenesis and restores proper granule cell integration. (A, B) Representative confocal images of DCX-TOM□ granule cells in the dorsal (A) and ventral (B) DG across experimental groups. Scale bars: 80□μm (A), 200□μm (B). DCX□ newborn granule cells (magenta) were labeled by tamoxifen-induced recombination in DCXCreERT2::TdTOM^lox/lox^ mice. (C) Quantification of ventral DCX-TOM□ cells revealed significantly increased neurogenesis in DMT- and DMT-d28–treated mice compared to Non-UCMS controls, with both groups also surpassing DMT+Iso and fluoxetine. (D) In the dorsal DG, UCMS-saline animals showed reduced neurogenesis compared to Non-UCMS controls. All treatments led to significantly increased neurogenesis compared to UCMS-saline, with DMT-treated animals showing the strongest response. (E) Representative image of ectopic DCX-TOM□ cells in a UCMS-saline animal. White arrows indicate DCX□ cells located outside the granule cell layer. Scale bar: 100□μm. (F) Quantification of ectopic DCX-TOM□ cells. UCMS-saline animals exhibited a marked increase in ectopic cells compared to Non-UCMS, indicating disrupted neuronal integration. All treatments significantly reduced ectopic neurogenesis relative to UCMS-saline. All data are presented as individual values with mean ± SEM. Analyses were performed using one-way ANOVA followed by Tukey’s post-hoc test. Sample sizes: Non-UCMS (n = 7), all other groups (n = 8).

Additionally, analyses within the dorsal hippocampus (Figure 4D) revealed that UCMS-saline animals showed reduced neurogenesis compared to Non-UCMS controls (p = 0.0044), indicating that chronic stress suppressed neurogenic activity in this region. Moreover, all treatments led to markedly increased neurogenesis relative to UCMS-saline animals, including DMT (p < 0.0001), DMT+Iso (p = 0.0002), fluoxetine (p < 0.0001), and DMT-d28 (p < 0.0001). DMT-treated mice exhibited the strongest neurogenic response and were the only group to significantly exceed Non-UCMS levels (p = 0.0060), suggesting not only a reversal of stress-induced deficits but a net increase in neurogenesis. DMT-treated animals also differed significantly from those given DMT under isoflurane (p = 0.0482), pointing to a possible role of wake psychedelic experience in neurogenesis. Fluoxetine and DMT-d28 groups showed intermediate DCX-TOM□ cell counts, although neither differed significantly from the DMT group (p = 0.5197 and p = 0.0892, respectively), suggesting a moderate, yet measurable, effect of these treatments on dorsal neurogenesis.

We also examined the incidence of ectopic DCX-TOM□ cells, indicative of disrupted neuronal integration due to stress (Figure 4E, F). Saline-treated UCMS mice displayed a significant increase in ectopic cell numbers compared to Non-UCMS controls (p < 0.0001), reflecting aberrant neurogenesis. Strikingly, all treatment groups showed significantly reduced ectopic cell counts relative to UCMS-saline, including DMT (p < 0.0001), DMT+Iso (p < 0.0001), fluoxetine (p = 0.0003), and DMT-d28 (p < 0.0001). Among these, DMT and DMT+Iso reduced ectopic cell incidence to levels statistically indistinguishable from Non-UCMS animals (p > 0.9999 and p = 0.9979, respectively), suggesting near-complete normalization. In contrast, fluoxetine- and DMT-d28-treated mice did not fully normalize ectopic integration and still differed significantly from Non-UCMS (p < 0.0001 and p = 0.4223, respectively). Furthermore, both DMT and DMT+Iso outperformed fluoxetine in reducing ectopic neurogenesis (p < 0.0001 and p = 0.0002), and fluoxetine in turn was significantly more effective than DMT-d28 (p = 0.0140). No significant differences were observed between the DMT and DMT+Iso groups (p = 0.9998), nor between either of those and DMT-d28 (p = 0.4949 and p = 0.6616). These findings highlight a superior efficacy of DMT-based interventions, particularly when administered after UCMS, in restoring proper neuronal integration patterns.

## Discussion

This study presents, to our knowledge, the first preclinical investigation into the antidepressant potential of a single dose of DMT using the UCMS paradigm in mice, a validated rodent model that mimics core features of MDD, including anhedonia, behavioral despair, and cognitive impairments [43, 45]. In doing so, we address several open questions concerning DMT’s capacity to reverse or prevent stress-induced depression-like phenotypes, its impact on adult hippocampal neurogenesis, and the necessity of the conscious psychedelic experience for its therapeutic effects. Our model successfully induced a robust depressive-like state, as evidenced by classic phenotypic hallmarks, such as reduced sucrose preference, increased immobility in the tail suspension test, and impaired pattern separation in the delayed non-match to place. These alterations were accompanied by reduced abGC production and abnormal ectopic integration of newborn neurons, consistent with previous findings linking stress to hippocampal disorganization and neurogenic disruption [3, 43, 45].

To date, most animal studies on psychedelics have emphasized anxiety-like phenotypes, with limited focus on depression [37, 46], likely due to the logistical ease of rapid, innate-behavior-based assays that are relatively quick to set up, require minimal training, and yield clear, measurable outcomes [43, 47]. In contrast, depression models, such as the UCMS paradigm, often involve prolonged exposure to stressors over weeks to induce depressive-like behaviors [48, 49]. These models require careful standardization of stress protocols and more extensive behavioral assessments, such as sucrose preference tests for anhedonia or cognitive tasks for memory impairments. Consequently, research on depressive conditions, which often involve distinct neurobiological mechanisms and behavioral manifestations, has lagged behind. While anxiety models have provided valuable insights into the therapeutic potential of psychedelics, they do not fully address the complexities of depression.

The present work helps bridge this gap by applying a clinically relevant and ethologically robust depression model to evaluate DMT’s effects on anhedonia, despair, cognition, and neurogenesis. We tested three DMT treatment regimens: a single post-stress dose (DMT), an early-dose administered midway through stress (DMT-d28), and a dose delivered under isoflurane anesthesia (DMT+Iso), to evaluate the timing and experiential contributions of treatment. Chronic fluoxetine served as a positive control. The findings reveal distinct effects depending on the timing of DMT administration and both surpass classical antidepressant treatment.

Administration of DMT after UCMS reversed all core depressive-like phenotypes. Treated mice exhibited restored sucrose preference, reduced immobility in the TST, and recovered pattern separation ability. These behavioral improvements were accompanied by significantly increased abGC production in both the dorsal and ventral DG, exceeding non-stressed levels in the dorsal DG. Furthermore, DMT normalized the rate of ectopic abGC integration, contrasting with fluoxetine’s partial effects. These findings echo and extend previous reports that serotonergic psychedelics can induce rapid and sustained antidepressant-like effects in rodent chronic stress models, including UCMS [50]. Notably, our data align with these studies not only behaviorally, but also at the level of hippocampal circuit repair, adding to the growing evidence that neurogenesis is a key mediator of psychedelic-induced resilience.

DMT administration during UCMS (DMT-d28) offered partial protection gainst the development of depressive-like behaviours. These mice exhibited preserved sucrose preference and reduced immobility in the TST, but failed to recover cognitive performance in the DNMP task. Histologically, neurogenesis was enhanced relative to untreated UCMS animals and ectopic integration was reduced, to a lesser extent but statistically similar to the post-stress DMT group. However, this early DMT administration did not rescue the UCMS-induced cognitive deficits. These findings suggest that early psychedelic intervention may confer prophylactic benefits against stress-induced emotional deficits but may be insufficient to fully counteract stress-induced hippocampal dysfunction without subsequent behavioral consequences. This is consistent with prior findings that psychedelic effects on neuronal plasticity lasted for several days after treatment, particularly in domains like learning and cognitive flexibility [51], although the extent of these changes may be limited by the post-treatment environment.

A key question in psychedelic research concerns the necessity of subjective psychedelic experiences for therapeutic outcomes [44]. By administering DMT under general anesthesia, we dissociated its physiological effects from its conscious phenomenology. Remarkably, DMT+Iso animals showed antidepressant-like outcomes and neurogenic enhancements comparable to the fully awake DMT group. This supports a mechanistic model in which DMT’s antidepressant actions are mediated by downstream neuroplastic cascades rather than by the subjective psychedelic experience itself. These findings are consistent with recent reports that psilocybin and LSD retain therapeutic efficacy in rodents even when 5-HT2A receptor signaling is pharmacologically blocked, provided neurotrophic signaling remains intact [50, 51].

Our work also highlights the role of adult hippocampal neurogenesis [24] in the antidepressant effects of DMT. Both DMT administered after and during stress significantly increased abGC production. The reduction in ectopic abGCs, which are indicative of disrupted hippocampal circuitry in stress, further supports the idea that DMT’s therapeutic effects involve the restoration of proper neuronal integration. Compared to fluoxetine, DMT produced stronger effects on both neurogenesis and behavior, highlighting its rapid and possibly more targeted mechanism of action. Together with previous results using 5-MeO-DMT [40], as well as harmala alkaloids like harmine and tetrahydroharmine which also promote neurogenesis [24, 52], our findings suggest that multiple components of the *ayahuasca* brew may converge on shared neurogenic and plasticity-enhancing pathways. These synergistic effects support a polypharmacological framework for psychedelic-induced circuit repair.

Moreover, the observation that post-stress DMT restored pattern separation deficits suggests a role for DMT in promoting hippocampal-dependent cognitive flexibility. Notably, chronic fluoxetine treatment achieved comparable improvements in the DNMP task, indicating that both treatments were effective in reversing UCMS-induced cognitive impairments. However, the behavioral and neurogenic profiles of the two interventions diverged. While fluoxetine failed to significantly restore sucrose preference, DMT robustly reversed anhedonia, suggesting a more pronounced effect on motivational and reward-related processes. At the neurobiological level, fluoxetine did not significantly enhance abGC production in the ventral hippocampus and only partially reduced ectopic cell migration. In contrast, DMT markedly increased neurogenesis across both hippocampal poles and nearly normalized aberrant integration. These findings suggest that DMT may involve not only 5-HT2A but also 5-HT2B and 5-HT2C receptors, both of which modulate plasticity and neurogenesis [52] or other neurotrophic signaling [51]. Whether these effects reflect direct receptor actions, indirect neurotrophic mechanisms, or a combination thereof remains to be clarified in future work.

In conclusion, this study provides preclinical evidence that a single dose of DMT can robustly reverse depression-like behaviors induced by chronic stress and enhance adult hippocampal neurogenesis. These effects appear independent of the conscious psychedelic state and stronger than those achieved by chronic fluoxetine. Together, our findings support the development of DMT as a rapid-acting antidepressant targeting circuit-level restoration. Moreover, our results show the importance of administration timing and behavioral context in shaping therapeutic outcomes. Given its rapid onset, sustained effects, and broad impact on both affective and cognitive domains, DMT emerges as a promising candidate for treating depression, particularly in individuals who do not respond to traditional monoaminergic therapies. Further investigation into optimal dosing, safety, and mechanisms will be essential to translate these findings into clinical strategies for fast-acting and scalable interventions.

## Methods

### Drugs

DMT (N,N-Dimethyltryptamine) was extracted and purified in-house from Mimosa tenuiflora root bark. The DMT free base was isolated and purified to an average purity of 99.0 ± 0.4% (range: 98.4–99.5%) as confirmed by gas chromatography-mass spectrometry (GC-MS). The compound was stored in dimethyl sulfoxide (DMSO) at *−*20 °C and diluted fresh in saline (1:9, DMSO:saline) on the day of treatment to reach 30 mg/kg dose. Fluoxetine was procured in pure form from a local compounding pharmacy (Ao Pharmacêutico) and diluted in saline to 20 mg/kg dose prepared fresh twice a week. Tamoxifen (Sigma-Aldrich, T5648) was prepared in autoclaved sesame oil (Sigma-Aldrich, S3547) in 100 µg/kg and stored at 4º C for no longer than five days.

### Animals and treatment

A total of 48 male DCXCreERT2::TdTOMlox/lox mice with a C57BL/6 genetic background, aged postnatal day 45 to 70, were used in this study. The animals were obtained from a local biotherium and pseudo-randomly assigned to six experimental groups, ensuring that littermates were distributed across treatments to mitigate litter-specific effects. All mice were housed under standard conditions, including a 12-hour light/dark cycle, with food and water available ad libitum. Experiments were conducted in six batches of eight animals each due to the extensive nature of the UCMS protocol. Each batch included at least one representative from each treatment group to maintain consistency across experimental conditions. The experimental design is illustrated in Figure 1A. For the Sucrose Preference Test (SPT) and the Delayed Non-Matching to Place (DNMP) task, environmental parameters such as food, water, and lighting were adjusted to align with the respective testing protocols 44–46.

Mice in all DMT-treated groups received a single intraperitoneal injection of 30mg/kg DMT with a final volume of animal weight (in grams) x 10µl (i.e, 20g animals received 200 µl) and were then returned to their home cages, where food and water were provided ad libitum. For the group treated during the ongoing UCMS protocol (DMT28d), the stressor was applied five hours prior to DMT administration to ensure a lower-stress baseline at the time of treatment, aligning conditions with other experimental groups. For the DMT-Iso group, mice were anesthetized with inhaled isoflurane before receiving DMT. Isoflurane anesthesia was initiated at a concentration of 5% for induction and maintained at 1.5–3% with a flow rate of 0.8 L/min. Anesthesia began 30 minutes prior to the DMT injection and continued for an additional 90 minutes after the treatment to ensure consistent exposure.

Fluoxetine-treated mice were administered fluoxetine intraperitoneally at a dose of 20 mg/kg daily by an experienced caretaker, starting on postnatal day 42 and continuing for 30 days. Treatment concluded one day before the initiation of behavioral testing to ensure the absence of residual drug effects during testing. Tamoxifen was administered to all groups at a dose of 100 µg/kg intraperitoneally for three consecutive days, beginning on postnatal day 48. This timing was carefully chosen to induce recombination in DCXCreERT2::TdTOMlox/lox mice, specifically labeling cells expected to integrate into hippocampal network during behavioral testing [40]. On the day of tamoxifen treatment, mice were housed under the same conditions to minimize variability.

### UCMS protocol and confirmation of depression-like symptoms onset

The UCMS protocol is a chronic stress paradigm designed to induce depression-like behaviors through prolonged exposure to unpredictable stressors. All mice, except those in the Non-UCMS group, were subjected to the UCMS regimen. Non-UCMS mice underwent gentle handling during the same period as the stressors. The UCMS protocol was implemented with minimal modifications as described by Monteiro et al., 2015 [53], and detailed in Supp. Table 1. A single aversive stressor was applied daily over an 8-week period. The stressors included confinement, where mice were placed in 50 ml plastic tubes (Falcon) with openings at both ends for 90 minutes; agitation, involving groups of five mice placed in a plastic box on an orbital shaker set at 150 rpm for 90 minutes; exposure to rat feces, a natural predator’s scent, where mice were subjected to 60 grams of soiled rat bedding for 90 minutes followed by cage cleaning; exposure to hot air jets from an 800 W hairdryer for 8 minutes, with minimal noise, while the box temperature was maintained at 35–40 ºC, monitored by a digital thermometer; nighttime illumination, in which mice were exposed to light for 2 hours during the dark phase; a reversed light/dark cycle for two consecutive days; and an inclined cage condition, where cages were tilted at a 45º angle for 90 minutes.

The stressors were presented unpredictably, as exemplified in Supp. Table 1, with the order randomized for the first experimental batch and then maintained consistently for subsequent batches. To confirm the development of a depressive-like phenotype, body weight and coat condition were monitored weekly, as shown in Figure 1 B-D. In addition, all animals underwent the SPT at three time points: pre-UCMS, post-UCMS, and post-pharmacological treatment. Animals with a sucrose preference ratio over water of less than 60% at baseline were excluded (n=1). Similarly, animals with a sucrose preference ratio over water of more than 75% after UCMS were excluded (n=2).

### Behavioral Experiments and Analysis

Following UCMS exposure and treatment completion, behavioral tests were conducted to individually assess anhedonia, apathy, cognition, and anxiety. Behavioral scoring and assessment was performed by experienced observers (R.V.L.dC., T.S., or T.C.M.) and computational tools. The first tool was DeepLabCut (DLC), a deep-learning-based software that has shown reliable results assessing animal behavior [54]. The second tool was AnimalTA, a tracking software that allows fast processing of a high number of multi-arena videos [55]. Custom Python or MATLAB codes were developed for downstream analysis of tracking outputs and are available upon request. The selected tests were performed as follows:

The Open Field Test (OFT) consisted in placing animals in an open arena with dimensions of 40 cm (length) × 32 cm (width) × 15 cm (height) for 10 minutes to assess general locomotor activity and anxiety-related behavior. AnimalTA [55] was used for tracking, and the trajectories were analyzed using our custom MATLAB pipeline.

Anxiety was evaluated by the elevated plus maze test (EPM). Animals were placed in a cross-shaped maze elevated 52 cm above the ground. The maze consisted of four arms, each 62 cm (length) × 6.5 cm (width), with two opposing arms enclosed by 20 cm high walls and two open arms. Each animal was tested for 10 minutes. AnimalTA was used to extract movement trajectories, which were then analyzed using the same custom MATLAB codes described above. Additionally, we manually analyzed the number of head dips outside the arena by a trained researcher.

The Sucrose Preference Test (SPT) assessed anhedonia, where mice were individually placed in their cages and habituated to drinking water from two bottles for 24 hours prior to the test. One of the bottles was then replaced with a 1% sucrose solution, and after 24 hours, the bottle positions were switched to eliminate location bias. Consumed volume was assessed by an experienced researcher.

The Tail Suspension Test (TST) evaluated behavioral despair. Animals were suspended by their tails 45 cm above the ground in a 60 cm (height) × 15 cm (depth) × 70 cm (width) square box. Their tails were gently secured with adhesive tape to a wooden shaft at a height of 45 cm. A small climb blocker, made from a 4 cm long cylinder cut from a 15 ml Falcon tube, was placed around their tails to prevent tail-climbing behavior, which is commonly observed in C57BL/6 mice. Since this test requires frame-by-frame pose estimation, we used DLC for tracking and the outputs were analyzed our custom Python script.

The Radial Arm Maze Test (RAM) assessed cognitive function, specifically reference and working memory. The maze consisted of eight arms, each 30 cm (length) × 8 cm (width) × 30 cm (height), connected to a central round platform with a 20 cm diameter. Each arm was equipped with removable doors and small wells at the end for food rewards, which were kept out of sight. The Delayed Non-Matching to Place (DNMP) paradigm was employed, consisting of sample and test phases. During the sample phase, the mouse was confined in the start arm without a reward while a single reward (a loop of Fruit Loops™) was placed in a designated sample arm located two arms away. Other arms remained closed, and the animal was allowed to retrieve the reward within four minutes. During the test phase, the mouse was returned to the start arm, and a reward was placed in a new arm. The mouse had to distinguish the previously visited sample arm from the newly opened reward arm to locate the food. A correct first attempt was recorded as a success, while errors were allowed to be corrected within 10 minutes, though errors were still noted. To minimize reliance on olfactory cues, the maze was rotated between phases (∼20 seconds), and the apparatus was thoroughly cleaned between subjects. The location of the reward arm was rotated relative to the sample arm by a factor of S# (where S represents the separation and # denotes the number of arms separating the two open arms). The test was repeated 20 times per condition (4 trials/day over 15 days) for each separation condition (S2, S3, and S4), resulting in 60 trials per condition per subject, as exemplified in Figure 3A. Animals with deficits in pattern separation were expected to show difficulty specifically in the higher difficulty S2 condition. To ensure task motivation, animals had limited access to food during the 15 days of testing and were habituated to the reward five days prior to maze training. All data was recorded manually during the experiment by an experienced researcher.

### Tissue Processing and Histological Analysis

Following the completion of behavioral experiments, animals were euthanized on day 78 via an overdose of Ketamine Hydrochloride (150 mg/kg) combined with Xylazine (8 mg/kg). They were then positioned in dorsal decubitus and perfused transcardially with 30 mL of phosphate-buffered saline (PBS), followed by 30 mL of 4% paraformaldehyde (PFA). After perfusion, craniotomies were performed to extract the brains, which were then submerged in 4% PFA at 2–8 °C overnight for fixation. The following day, brains were cryoprotected by immersion in 30% sucrose solution for 24 hours. Once cryoprotected, they were briefly rinsed in 0.1M phosphate buffer (PB), rapidly frozen by immersion in -80 °C isopentane at, and then stored at a -80 °C freezer before further processing.

Frozen brains were horizontally sectioned into 25 µm thick slices using a cryostat. Sections were collected on gelatin-coated slides in an alternating series, with a 200 µm spacing between slices, covering the entire hippocampal region (AP 2.6 - 4.4 mm, according to Paxinos, 2012). Tissue sections were mounted with Fluoromount-G™ medium containing DAPI (ThermoFisher, cat. 00-4959-52) and examined under a ZEISS reflected light microscope. Stereoinvestigator software (MBF Bioscience) was used to assist in cell quantification. tdTOM+ cells were manually counted in both hippocampi by an experimenter blinded to the experimental groups. Ectopic cells were defined as tdTOM+ cells located >20 μm away from the Granule Cell Layer (GCL) in any direction.

## Statistical analysis

All data were tested for normality using D’Agostino and Pearson’s omnibus normality test and were confirmed to follow a normal distribution. The sample size varies across some tests due to technical errors during behavioral recordings or equipment failures, which resulted in the loss of specific trials while the affected animals remained in the study for all other assessments. Data are expressed as Mean ± SEM, unless stated otherwise. Statistical significance was set at p < 0.05. Group comparisons were conducted using one-way ANOVA followed by Tukey’s multiple comparisons test or two-way ANOVA followed by Holm-Sidak’s multiple comparisons test. All statistical analyses were performed using GraphPad Prism 10 (San Diego, CA, USA) and .csv processing software.

## Ethical Considerations

All procedures were conducted in accordance with the ethical guidelines set by the Brazilian National Council for the Control of Animal Experimentation (CONCEA). The protocols were approved by the institutional animal care and use committee of the Federal University of Rio Grande do Norte (approval numbers 036/2019 and 186.036/2019).

## Figure Legends

**Supplementary Figure 1. Anxiety-like phenotypes were assessed by open field and elevated plus maze tests**. (A) Representative illustration of movement tracking paths (top panel) and occupancy density plots (bottom panel) from the open field test of a representative mouse from the UCMS DMT group. (B) Quantification of total distance traveled (cm) in the open field test. (C) Percentage of time spent in the center area relative to total test duration in the open field test. (D) Representative illustration of movement tracking paths in the elevated plus maze from a representative mouse of the UCMS DMT group. (E) Percentage of total test time spent in the open arms of the elevated plus maze. (F) Number of head dips as an indicator of exploratory behavior in the elevated plus maze. All data are presented as individual values with mean ± SEM. Statistical analyses were conducted using one-way ANOVA followed by Tukey’s post-hoc test. Sample sizes for open field test: Non-UCMS (n=8), UCMS Saline (n=5), UCMS Fluox (n=4), UCMS DMT (n=6), UCMS DMT+ISO (n=7), UCMS DMTd28 (n=6). Sample sizes for elevated plus maze test: Non-UCMS (n=8), UCMS Saline (n=5), UCMS Fluox (n=4), UCMS DMT (n=6), UCMS DMT+ISO (n=7), UCMS DMTd28 (n=6).

