## Supplementary Table 1 for "Single-dose DMT reverses anhedonia and cognitive deficits via restoration of neurogenesis in a stress-induced depression model"

| | Monday | Tuesday | Wednesday | Thursday | Friday | Saturday | Sunday |
|---|---|---|---|---|---|---|---|
| **1st week** | BW; CS; Hot air jet | Shaking | Confinement | Rat feces | Confinement | Confinement | Tilted cage |
| **2nd week** | BW; CS Confinement | Shaking | Rat feces | Confinement | Shaking | Rat feces | Confinement |
| **3rd week** | BW; CS Confinement | Rat Feces | Confinement | Hot air jet | Rat feces | Confinement | Shaking |
| **4th week** | BW; CS Rat feces | Confinement | Rat feces | Shaking | Confinement | Shaking | Hot air Jet |
| **5th week** | BW; CS Confinement | Tilted cage | Hot air jet | Confinement | Rat feces | Interted Light cycle | Rat feces |
| **6th week** | BW; CS Hot air Jet | Shaking | Confinement | Tilted Cage | Nighttime illumination | Confinement | Hot air jet |
| **7th week** | BW; CS Confinement | Rat feces | Confinement | Hot air jet | Shaking | Confinement | Nighttime illumination |
| **8th week** | BW; CS Shaking | Confinement | Rat feces | Shaking | Confinement | Confinement | Shaking |
| **9th week** | BW; CS | | | | | | |

**Supplementary Table 1:** Chronogram for the UCMS protocol. B.W.: Body Weight; C.S.: Coat Score.
