## Supplementary figures and images for "Single-dose DMT reverses anhedonia and cognitive deficits via restoration of neurogenesis in a stress-induced depression model"

### Supplementary Figure 1

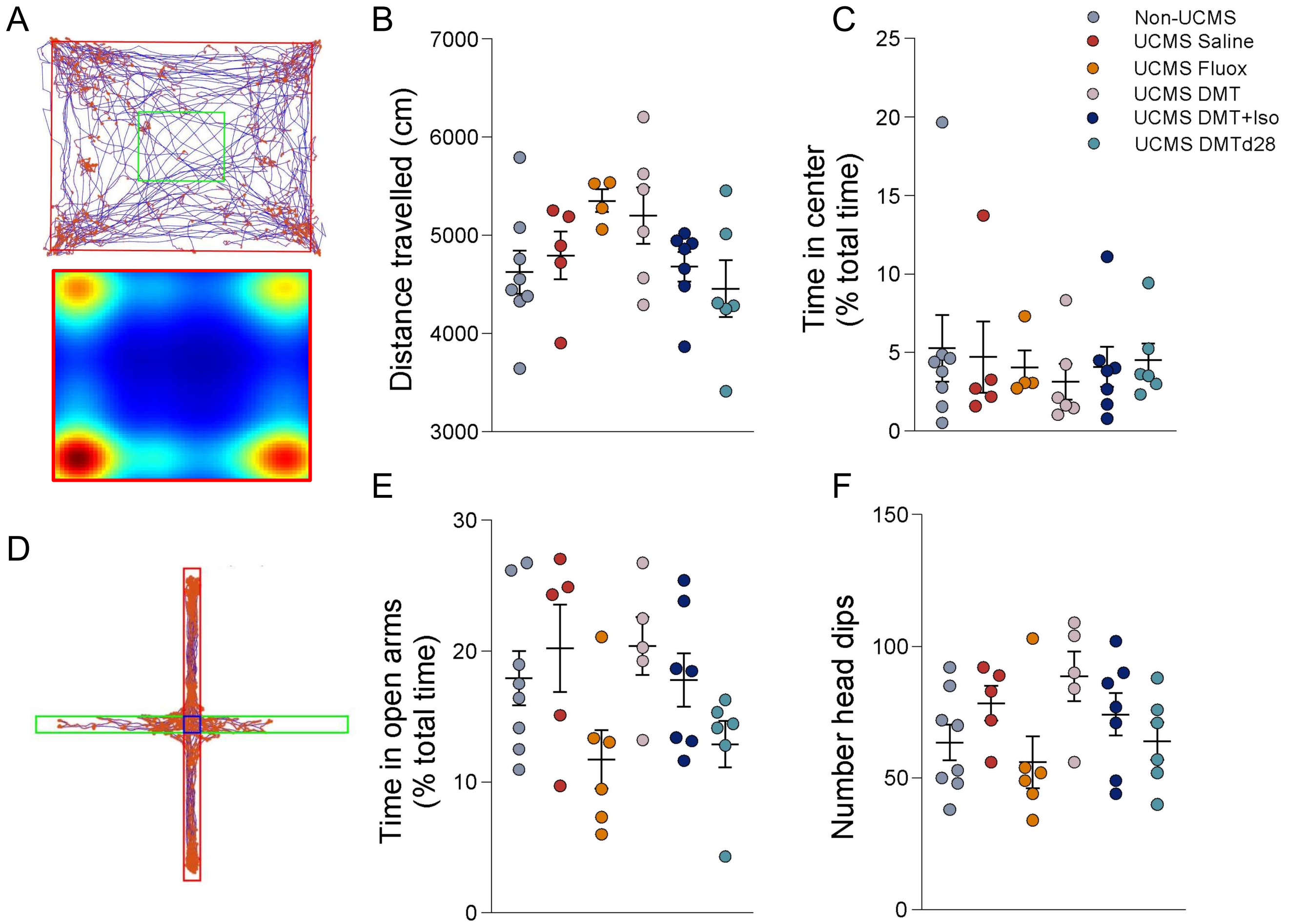


Supplementary Figure 1
